# Contrasting mechanisms for hidden hearing loss: synaptopathy vs myelin defects

**DOI:** 10.1101/2020.10.04.324335

**Authors:** Maral Budak, Karl Grosh, Gabriel Corfas, Michal Zochowski, Victoria Booth

**Author notes:** Corresponding authors (VB), (GC), (MZ). These authors contributed equally to this work (Michal Zochowski, Victoria Booth).

## Abstract

Hidden hearing loss (HHL) is an auditory neuropathy characterized by normal hearing thresholds but reduced amplitude of the sound-evoked auditory nerve compound action potential (CAP). It has been proposed that in humans HHL leads to speech discrimination and intelligibility deficits, particularly in noisy environments. Animal models originally indicated that HHL can be caused by moderate noise exposures or aging, and that loss of inner hair cell (IHC) synapses could be its cause. A recent study provided evidence that transient loss of cochlear Schwann cells also causes permanent auditory deficits in mice which have characteristics of HHL. Histological analysis of the cochlea after auditory nerve remyelination showed a permanent disruption of the myelination patterns at the heminode of type I spiral ganglion neuron (SGN) peripheral terminals, suggesting that this defect could be contributing to HHL. To shed light on the mechanisms of different HHL scenarios and to test their impact on type I SGN activity, we constructed a reduced biophysical model for a population of SGN peripheral axons. We found that the amplitudes of simulated sound-evoked SGN CAPs are lower and have greater latencies when the heminodes are disorganized, i.e. they are placed at different distances from the hair cell rather than at the same distance as seen in the normal cochlea. Thus, our model confirms that disruption of the position of the heminode causes desynchronization of SGN spikes leading to a loss of temporal resolution and reduction of the sound-evoked SGN CAP. We also simulated synaptopathy by removing high threshold IHC-SGN synapses and found that the amplitude of simulated sound-evoked SGN CAPs decreases while latencies remain unchanged, corresponding to what has been observed in noise exposed animals. This model can be used to further study the effects of synaptopathy or demyelination on auditory function.

**Author summary:** Hidden hearing loss is an auditory disorder caused by noise exposure, aging or peripheral neuropathy which is estimated to affect 12-15% of the world’s population. It is a ‘hidden’ disorder because subjects have normal hearing thresholds, i.e., the condition cannot be revealed by standard audiological tests, but they report difficulties in understanding speech in noisy environments. Studies on animal models suggest two possible pathogenic mechanisms for hidden hearing loss: (1) loss of synapses between inner hair cells and auditory nerve fibers, and (2) disruption of auditory-nerve myelin. In this study, we constructed a computational model of sound-evoked auditory neuron fiber activity and auditory nerve compound action potential to understand how each one of these mechanisms affects nerve transmission. We show that disruption of auditory-nerve myelin desynchronizes sound-evoked auditory neuron spiking, decreasing the amplitude and increasing the latency of the compound action potential. In addition, elongation of the initial axon segment may cause spike generation failure leading to decreased spiking probability. In contrast, the effect of synapse loss is only to decrease the probability of firing, thus reducing the compound action potential amplitude without disturbing its latency. This model, which accurately represents the in vivo findings, could be useful to make further predictions on the consequences of HHL and extend it to explore the impact of synaptopathy and myelinopathy on hearing.

## Introduction

Hidden hearing loss (HHL) is defined as an auditory neuropathy characterized by changes in neural sound-evoked output of the auditory nerve (AN) without hearing threshold elevation [1]. The prevalence of HHL has been estimated at 12-15% based on recent surveys where subjects with normal hearing thresholds reported difficulties in hearing, especially in noisy environments [2, 3]. HHL has been detected in animal models and humans by measuring the neural responses to suprathreshold sound via tests, such as auditory brainstem response (ABR), a far-field response measured by head-mounted electrodes, or compound action potential (CAP), a near-field response measured from the round window. CAP and the first peak of ABR (ABR peak 1) represent the activity of type I spiral ganglion neurons (SGNs) in response to sounds [1].

There is mounting evidence that HHL can be caused by noise exposure, aging or peripheral myelin neuropathy [4–7]. After exposure to moderate noise, animals and humans have temporary shifts in auditory thresholds but permanent decreases in amplitude of ABR peak 1 [4–7]. Kujawa and Liberman (2009) showed that animals with this type of auditory pathology have a normal complement of hair cells and SGNs, but present with loss of a subset of synaptic connections between inner hair cells (IHCs) and SGNs. They also found that the degree of synapse loss correlates with the magnitude of the decrease in suprathreshold responses, supporting the idea that cochlear synaptopathy is the mechanism for noise-induced HHL [4]. Similar observations were made regarding aging, i.e. HHL and synapse loss are the first signs of age-related hearing loss and have the same time-course [5]. Importantly, it has been suggested that moderate noise and aging primarily affect synapses associated with high threshold/low spontaneous rate SGN fibers [8]. Since these fibers can respond to sound in high background noise even when the others have been saturated, their loss should lead to difficulties in processing speech in noisy environments [8].

Auditory processing requires proper myelination of auditory nerves [9]. Therefore, it has been hypothesized that peripheral neuropathy resulting from myelin disorders may be another cause of HHL. Individuals with peripheral neuropathies, such as Guillain-Barré Syndrome (GBS) [10] and Charcot-Marie-Tooth (CMT) disease [11] have been reported to have perceptual difficulties even when having normal auditory thresholds, indicating HHL. A recent study by Wan and Corfas (2017) showed that transient demyelination also causes HHL in mice, i.e. reduced ABR peak 1 amplitude with normal ABR thresholds [6]. In that study, acute demyelination was induced using genetically modified mice. This demyelination resulted in decreased ABR peak 1 amplitudes and increased ABR peak 1 latency without auditory threshold elevation or IHC-SGN synapse loss. Remarkably, these changes persisted even after remyelination of SGN fibers. Further investigation with immunostaining demonstrated that the organization of the heminodes, the nodal structures closest to the IHCs where action potentials are generated, were disrupted. These results suggested that the location of SGN heminodes is critical for normal auditory responses and that their disruption causes HHL.

In this study, we investigated the implications of these two HHL mechanisms, synaptopathy and myelinopathy, on sound-evoked spike generation and timing in SGNs. For this purpose, we constructed a reduced biophysical model consisting of a population of SGN fibers to investigate how synapse loss or disruption of myelin organization affect spike generation and transmission. Synaptopathy and myelinopathy were implemented by removing synapses and varying the position of SGN heminodes, respectively. Model results show that heminode disruption causes decay of the amplitude and increases the latency of sound-evoked CAPs. In addition, significant elongation of the initial axon segment causes spike generation failure leading to decreased spiking probability. In contrast, synaptopathy, solely decreases probability of firing, subsequently decreasing CAP peak amplitude without affecting its latency. These results are consistent with experimental observations [4, 6].

## Methods

### SGN fiber model

Type I SGNs are bipolar neurons with peripheral axon segments innervating IHCs and central axon segments projecting into cochlear nucleus (Fig 1A) [12]. In this study, a compartmental model of peripheral axons of type I SGNs was constructed using the NEURON simulator (version 7.6.2, [13]) as schematized in Figs 1B and C. For simplicity, we refer to peripheral axons of type I SGNs as SGN fibers, throughout the paper. Each fiber consists of an unmyelinated segment (length L_u_), a heminode (length L_h_) and 5 myelin sheaths following the heminode, separated by 4 nodes. Each compartment has passive membrane properties described by specific capacitance (C_m_) and specific membrane resistance (R_m_). Specific cytoplasmic resistance (R_a_) between each consecutive compartment was modified to obtain the speed of the action potential as 12-14m/s [14], based on the neural conduction velocity measurements of human auditory nerve [15]. Sodium and potassium channels were inserted along the SGN fibers, except the myelin sheaths, which only had passive membrane properties. The nominal conductances of both channel types at the unmyelinated segment was 15 times less than the nodes and the heminode [16], therefore action potential was initiated first at the heminode. The parameters for channel dynamics were taken from [14] (see S1 File), the stochastic channels in [14] were converted into deterministic ones for simplicity. This was done by multiplying channel density with the single ion channel conductance to obtain deterministic conductance values (see Table 1 for all parameters). The Nernst potentials for the ions Na^+^ (E_Na_) and K^+^ (E_K_) were set to 66 and −88 mV, respectively, and the resting potential (E_Rest_) was −78 mV [17]. Simulations were done at 37°C. The differential equations were solved by fully implicit backward Euler method with time step 5μs implemented in the NEURON simulation environment (see S1 File).

**Table 1:** Morphological, electrical and ion channel parameters of the different parts of a normal SGN fiber. Values as in [16] except for Ra and myelinated segment length which were modified for human SGN fibers.

| Parameters | Unmyelinated segment | Heminode | Myelin | Node |
| --- | --- | --- | --- | --- |
| Length ($\mu\text{m}$) | 10 | 1 | 40 | 1 |
| Diameter ($\mu\text{m}$) | 1.2 | 1.2 | 2.2 | 1.2 |
| $g_{\text{Na}}$ ($\text{S}/\text{cm}^2$) | 0.01208 | 0.1812 | 0 | 0.1812 |
| $g_{\text{K}}$ ($\text{S}/\text{cm}^2$) | 0.015 | 0.225 | 0 | 0.225 |
| $R_{\text{m}}$ ($\text{ohm}\cdot\text{cm}^2$) | 1662 | 1662 | 1300000 | 1662 |
| $C$ ($\mu\text{F}/\text{cm}^2$) | 0.05125 | 0.05125 | 0.0012 | 0.05125 |
| $R_a$ ($\text{ohm}\cdot\text{cm}$) | 8291.4 | | | |

$g_{\text{Na}}$ , maximal sodium conductance; $g_{\text{K}}$ , maximal potassium conductance; $R_{\text{m}}$ , specific membrane resistance; $C$ , specific capacitance; $R_a$ , specific cytoplasmic resistance.

**Fig 1.**
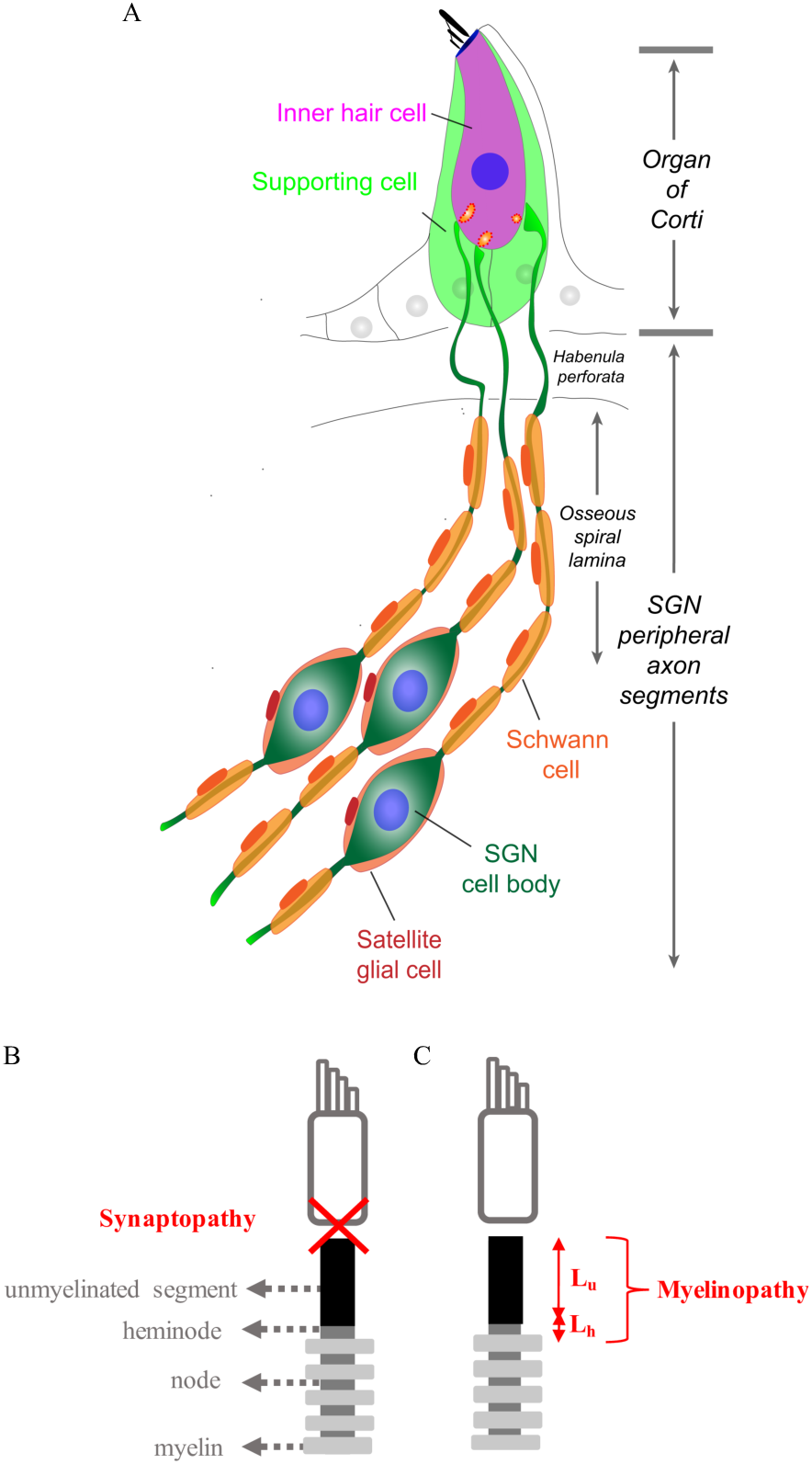
Diagram of model SGN fiber illustrating two mechanisms of hidden hearing loss. (A) Schematic illustration of type I SGNs, bipolar neurons innervating IHCs via myelinated peripheral projections. (B,C) Model peripheral fibers of type I SGNs (SGN fiber) consist of an unmyelinated segment at the peripheral end adjacent to the site of IHC synapses, followed by a heminode and 5 myelin sheaths with 4 nodes between them. Two mechanisms of hidden hearing loss are simulated: (B) synaptopathy, modeled by removing IHC-AN synapses, and (C) myelinopathy, modeled by varying the lengths of the unmyelinated segment (L_u_) or the heminode (L_h_).

### Sound representation

Increasing sound level increases the probability of neurotransmitter release from IHCs [18], therefore we defined the sound stimulus in terms of a release probability p_i_ at each IHC-SGN synapse (Fig 2A). Since type I SGNs also fire spontaneously [19], we set a release probability of p_spont_ in the absence of sound. To simulate the activity of SGNs in response to sound stimulus as described in [20], release probability was first increased sharply up to a defined peak (p_spont_+p_peak_), then allowed to decay due to adaptation to a constant level at the half-peak 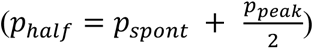 until the end of the sound stimulus. Thus, the release probability function p_i_(t) was defined as:

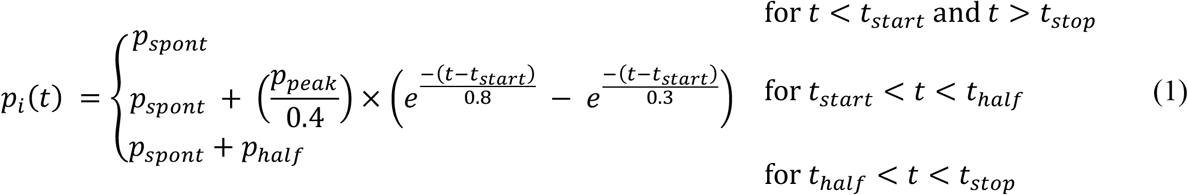

where t_half_ was the time at which the function decays to p_spont_+p_half_ after passing p_peak_, t_start_ and t_stop_ were the times when sound stimulus starts and ends, respectively.

**Fig 2.**
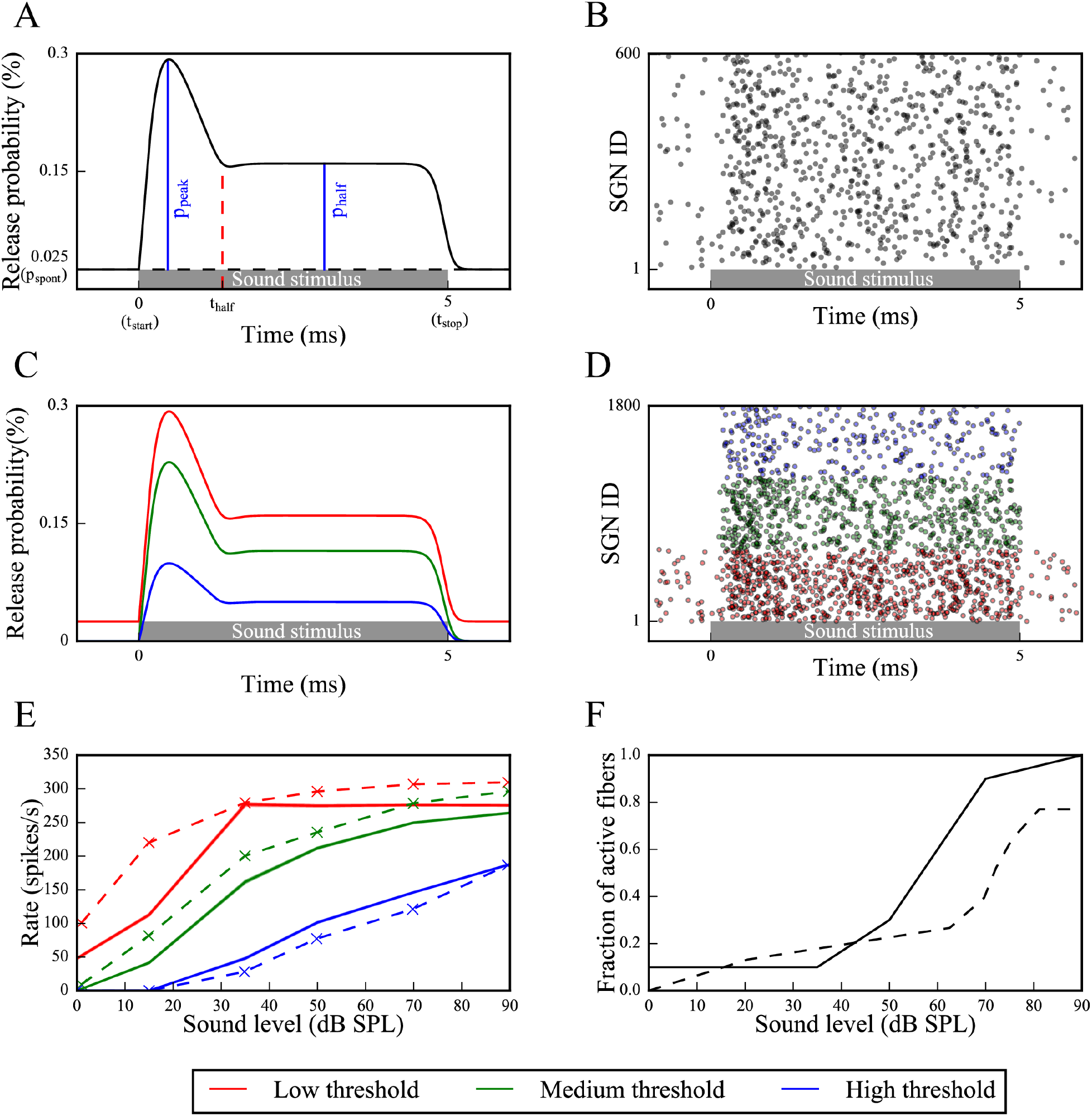
Sound-evoked activity of low, medium and high threshold SGN fibers results from increased vesicle release probabilities from corresponding IHC-SGN synapses. (A) Sound stimuli is modeled as an increased vesicle release probability from IHCs where release times are determined by a Poisson process (p_spont_: spontaneous release probability, p_peak_: maximum release probability, p_half_: release probability after adaptation, t_start_: sound start time, t_stop_: sound end time, t_half_: adaptation start time). For each release event, the corresponding SGN fiber is stimulated with a brief external current pulse, resulting in spiking activity; cumulative activity of an SGN fiber population is shown in (B), where gray dots represent spike times of each SGN fiber in a population defined as different SGN fiber IDs. (C) Three groups of SGN fibers, low (LT), medium (MT) and high (HT) threshold, were simulated based on their spontaneous firing rates and saturation profiles in response to sound, by defining p_peak_ and p_spont_ for each fiber type at each sound level. (D) Based on the release probabilities, different fiber types exhibit different cumulative responses (red dots: low threshold, green dots: medium threshold, blue dots: high threshold). Panels A-D are example simulations for simulated 50dB SPL. (E) The trend of spike rates of each fiber type for various sound levels in our model (solid lines) are comparable to experimental results (dashed lines) (Data taken from [19]). (F) For higher sound levels, due to the recruitment of SGN fibers with CFs near that of the simulated sound frequency, the fraction of activated SGN fibers increases for sound levels higher than 35dB SPL. At 90dB SPL, all 6000 fibers are activated (solid line: our model, dashed line: calculated based on spiral ganglion cell densities for different CFs [21] and IHC tuning curves [22]).

IHC-SGN synaptic release probability p_i_(t) was used to determine a Poisson process of IHC release that governed brief external stimuli to the corresponding nerve fiber to induce action potential generation. The external stimuli mimicking synaptic release from IHCs were simulated in the form of external current pulses with amplitude 0.024 nA and duration 0.05 ms (I_app_) applied at the beginning of the unmyelinated segment, unless otherwise stated (S3A Fig). The amplitude and duration of the stimuli were chosen to be close to the threshold stimulation needed to result in action potential generation at the heminode of putative control (L_u_=10 μm, L_h_=1 μm) fibers. The time of the action potential at the center of the heminode was taken as output (Fig 2B).

In our model, to reproduce experimentally observed shape of the stimulus mediated release [6], we used 5 ms long sound stimuli (*t_stop_* − *t_start_* = 5*ms*) and set the time from stimulus initiation to release probability reaching p_half_, i.e. (t_half_ − t_start_), to 1.4ms [20]. We varied p_spont_ and p_peak_ for different sound levels and fiber types (Table 2) to obtain experimental response properties [19].

**Table 2:** Spontaneous (p_spont_) and maximum (p_peak_) release probabilities for low- (LT), medium- (MT) and high-threshold SGN fibers (HT)

|  |  | LT | MT | HT |
| --- | --- | --- | --- | --- |
| $p_{\text{spont}}$ | | 0.00025 | 0 | 0 |
| $p_{\text{peak}}$ | 0 dB SPL | 0 | 0 | 0 |
|  | 15 dB SPL | 0.0007 | 0.0004 | 0 |
|  | 35 dB SPL | 0.0027 | 0.0017 | 0.00045 |
|  | 50 dB SPL | 0.0027 | 0.0023 | 0.001 |
|  | 70 dB SPL | 0.0027 | 0.0028 | 0.0015 |
|  | 90 dB SPL | 0.0027 | 0.003 | 0.002 |

$p_{\text{spont}}$ , spontaneous release probability; $p_{\text{peak}}$ , maximum release probability; LT, low-threshold SGN fiber; MT, medium-threshold SGN fiber; HT, high-threshold SGN fiber

### Defining different sound levels and fiber types

SGNs can be classified into 3 groups depending on their spontaneous firing properties, thresholds for sound-evoked activity and saturation profiles, namely low threshold (LT), medium threshold (MT) and high threshold (HT) fibers. Based on the measurements reported in [19], we modeled the properties of these three fiber groups as follows (Figs 2C-E): LT fibers have high spontaneous rates (18-100 spikes/s), low dynamic ranges, and reach their maximum discharge rate within approximately 30 dB sound pressure level (SPL). MT fibers have lower spontaneous firing (between 0.5 and 18 spikes/s), higher dynamic ranges, and show slower increase and saturation of spike rates with increasing SPL compared to LT fibers. HT fibers have very low spontaneous firing rates (<0.5 spikes/s), and response thresholds higher than ~20 dB SPL. For higher SPL, their spike rate increases linearly with sound intensity, therefore their dynamic range is the highest [19].

In our model, SPL is simulated by varying the peak probability, p_peak_, of the IHC-SGN synaptic release probability function. The fiber type response properties are obtained by defining spontaneous release probability, p_spont_, and scaling p_peak_ for each sound level. Figs 2C and 2D show IHC-SGN synaptic release probability functions and spike firing, respectively, for 70dB SPL for each fiber type (see Table 2 for all sound levels). For simplicity, we assumed MT and HT fibers do not fire spontaneously.

The intensity of sound stimuli affects the number of recruited type I SGNs as well. At lower sound levels, only the fibers with the characteristic frequencies (CFs) close to the stimulus fire. At higher SPL, the spatial profile of excitation spreads, and fibers with a broader range of CFs also respond to the stimulation. To introduce the recruitment of more fibers with increasing SPL, we considered the IHC tuning curves [22] and the density of SGNs based on their CF [21], and used 600 fibers for sound intensities of 35dB SPL and lower, 1800 fibers for 50 dB SPL, 5400 fibers for 70 dB SPL and 6000 for 90 dB SPL, with equal numbers of LT, MT and HT fibers (Fig 2F).

### Analyzing spike trains obtained from simulations

In response to simulated sound stimulus, each model SGN fiber fires a sequence of spikes (Fig 2D). We used three methods to analyze SGN fiber spike trains:

#### Measurement of time intervals between non-identical spike trains of SGN fiber populations

This metric, modified from a shuffled autocorrelogram measure in [23], was used to quantify temporal properties of SGN fiber spiking within a population based on the time intervals of the spikes between each non-identical pair of spike trains within the population. From all possible non-identical pairs of spike trains within a population, forward time intervals were measured between each spike *i* of the first spike train and spikes of the second spike train falling between the *i*-th and (*i+1)*-st spikes (Fig 3A). All time intervals from all pairs were tallied in a histogram and the histogram was reflected over y-axis, since each forward time interval of a pair (x,y) is a backward time interval of the pair (y,x).

**Fig 3.**
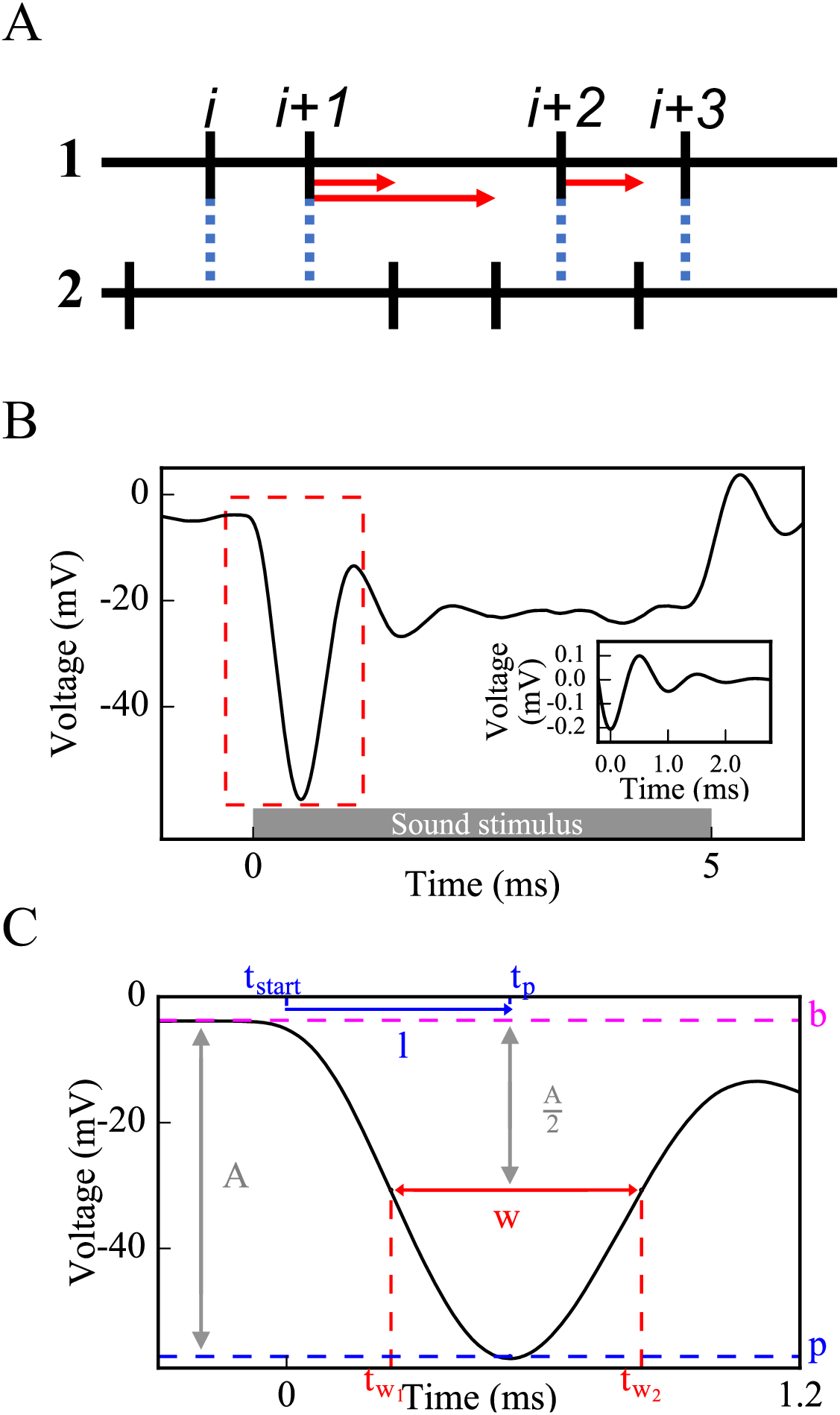
Methods used to evaluate cumulative activity of SGN fiber populations: pairwise spike time differences (A) and simulated CAP (B,C). (A) For each non-identical pair of spike trains from an SGN fiber population, forward time intervals are measured between each spike *i* of the spike train 1 and all spikes of the spike train 2 falling between *i* and *i+1*. Standard deviations of the distributions of these time intervals are calculated to evaluate synchronous spike timing in the SGN fiber population. (B) Each spike in Fig 2D is convolved with the unitary response of CAP [the inset of (B)] and convolutions from each spike are summed up to obtain a simulated CAP of the SGN fiber population. (C) Amplitude, latency and width are measured from the first peak of the simulated CAP [dashed rectangle in (B) is zoomed in for (C)] (b: baseline, p: peak, A: amplitude of the peak, t_p_: peak time, l: latency, w: width, t_w1_: half amplitude time before t_p_, t_w2_: half amplitude time after t_p_).

#### Convolution into the unitary response of compound action potential (CAP)

To yield a cumulative response of the activity of the population of SGN fibers and to be able to compare model results with in vivo ABR P1 results, we convolved each spike with the unitary response and summed them up to generate a population CAP (Fig 3B). In this study, we considered this computed CAP as equivalent to ABR P1. The unitary response U(t) was described as in [24]:

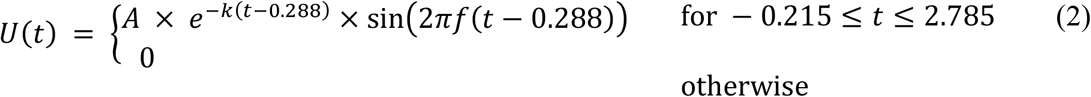

where *A* = 0.14*μV,k* = 1.44*ms*^−1^, *f* = 0.994*ms*^−1^ and t is the time (Fig 3B inset).

Fifty population CAPs were averaged to measure the width (w), amplitude (a) and latency (w) of the initial CAP peak more accurately, which were computed as:

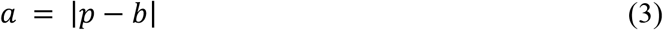

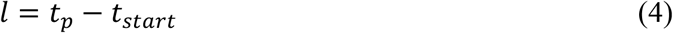

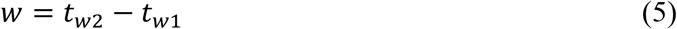

where p is the peak voltage, b is the baseline voltage, t_p_ is the time when the voltage equals p, and t_w1_ and t_w2_ are the times when the voltage equals 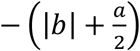 (the half-peak) before and after t_p_, respectively (Fig 3C).

#### Calculating spike probability and latency for each SGN fiber population

The probability that release events at IHC-SGN synapses resulted in spikes at the heminodes of an SGN fiber population was calculated by dividing the number of spikes at the heminode of each SGN fiber by the number of release events and averaging over all fibers within a population. Spike latency of an SGN fiber population was calculated by the time difference between a spike and a release preceding that spike averaged over all spikes of that population.

## Results

Using the model of the type I SGN fiber population, we investigated the effects of myelinopathy and synaptopathy on type I SGN spike generation and spike timing. We first simulated different myelinopathy scenarios by varying the length of the initial unmyelinated segment L_u_ (Fig 1C, from a putative control value of 10 μm) and the first heminode length L_h_ (from a control value of 1 μm) for all (i.e. LT, MT and HT) fibers. Next, we simulated synaptopathy by removing IHC-SGN synapses (Fig 1B) considering the cases where only synapses on HT fibers are affected or synapses on all fiber types are affected. Lastly, we investigated the combined effects of myelinopathy and synaptopathy.

### Effects of myelinopathy on SGN population activation patterns

Mouse studies have shown that transient demyelination and the subsequent remyelination alters the position of SGN heminodes, resulting in heminodes that are positioned farther from the IHC-SGN synapse and at variable positions, in contrast to healthy SGN fibers where heminodes on all fibers are aligned [6]. To identify the effect of this heterogeneity of heminode locations on SGN spike timing, we first considered a population of only LT fibers with different ranges of L_u_ values (Fig 4). Here, we denote 0% increase as the putative control fiber length (L_u_=10 μm), while 100% increase means L_u_ was varied between 10 and 20 μm across the population. We assessed the level of synchronization of spikes across the SGN fiber population by stimulating all fibers with the same 90dB simulated SPL with the identical IHC release pattern. As heterogeneity of L_u_ values was increased (Fig 4A), the population spike rate decreased reflecting spike generation failure on fibers with large L_u_. At the same time, variability in spike timing increased as illustrated in spike raster plots (Figs 4B, D, F, H show a portion of the generated spike trains, insets show timing of first spikes) and computed pairwise spike time intervals (Figs 4C, E, G, I, see Methods). These disruptions in spike generation and timing resulted in increased standard deviation of the distribution of pairwise spike time differences across the population (Fig 4A). These initial observations suggest that myelinopathy not only disrupts spike timing of SGNs within a population, but also leads to the loss of spikes.

**Fig 4.**
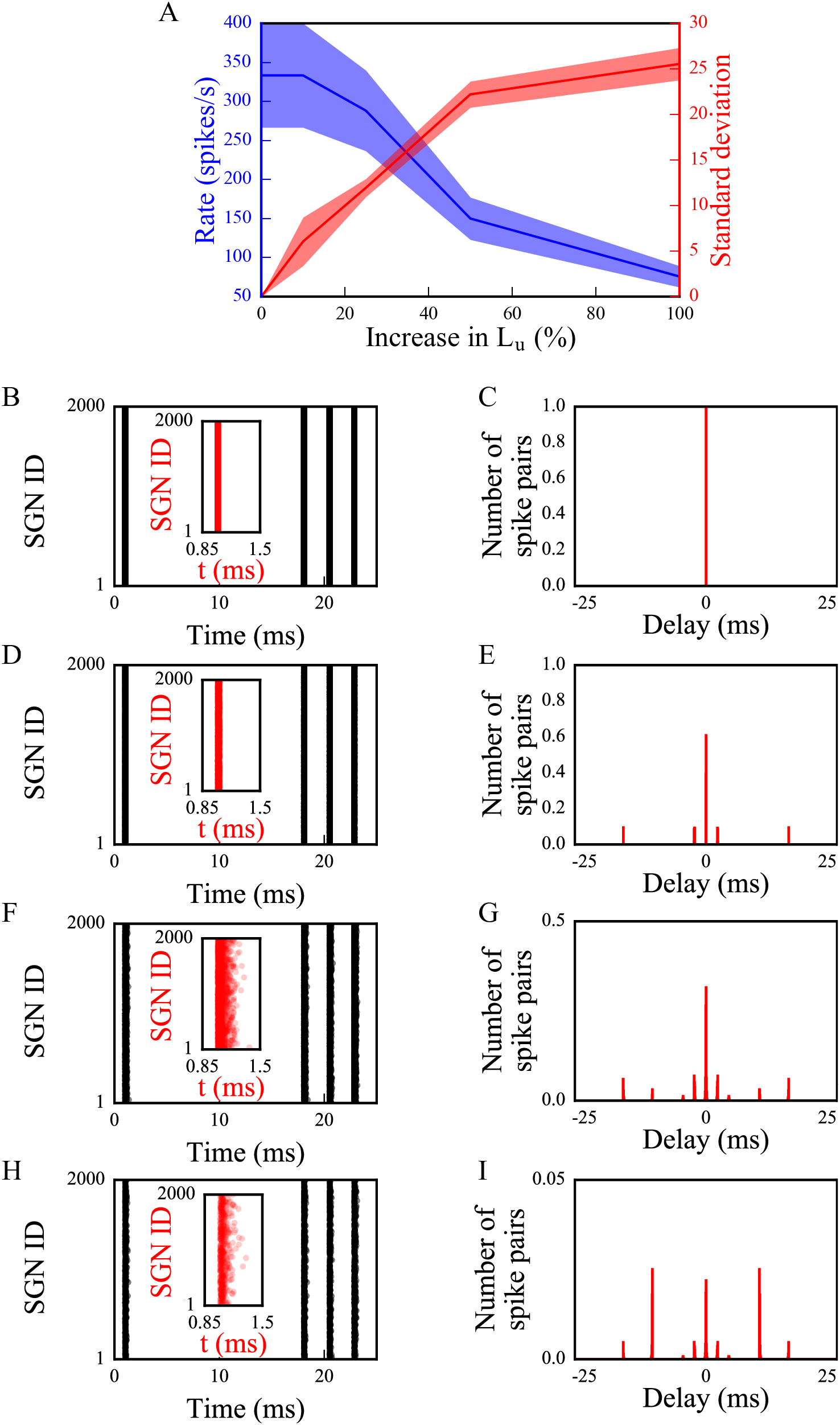
The synchronous activity of SGN fiber populations is disrupted and their response to sound is decreased with increasing levels of L_u_ heterogeneity. SGN fiber populations with different heterogeneity levels of L_u_ were stimulated using the identical simulated 90dB SPL stimulus. We assumed release events from all IHCs for the population occurred simultaneously. Raster plots of a portion of the generated spike trains [(B), (D), (F) and (H), insets: timings of first spikes] and corresponding histograms of time intervals between non-identical pairs of spike trains within a population [(C), (E), (G) and (I)] are shown for populations of SGN fibers with L_u_=10μm (0% increase in L_u_) [(B) and (C)], 10μm≤L_u_≤11μm (10% increase in L_u_) [(D) and (E)], 10μm≤L_u_≤12.5μm (25% increase in L_u_) [(F) and (G)] and 10μm≤ L_u_≤20μm (100% increase in L_u_) [(H) and (I)]. The ordinates of the histograms are normalized over the number of spike pairs with 0ms delay for the population where all fibers have L_u_=10μm (C). Simulations were done 3 times. Firing rate and standard deviations of time intervals are averaged for all populations in (A), shaded area represents the standard error of the mean.

To investigate effects of this disruption of spike generation and timing in the full model, CAPs were computed from spike responses of populations of LT, MT and HT SGN fibers subject to simulated myelinopathy. Responses of fiber populations with homogeneous initial unmyelinated segments (L_u_) or first heminode length (L_h_) values were investigated to see the gradual effect of variable myelination patterns on cumulative activity of SGN fibers. Additionally, populations with heterogeneous, random L_u_ or L_h_ values were simulated to represent a population heterogeneity induced by myelinopathy. We note that when increasing first heminode length (L_h_) the number of expressed channels (Na^+^ and K^+^) was kept constant consequently decreasing their density. However, when increasing initial unmyelinated segment length (L_u_), the density of expressed channels was kept constant consequently increasing their number. Results were not qualitatively different when these assumptions were reversed (see Discussion section). Model results show that, in response to a simulated 70 dB SPL stimulus, CAPs computed from SGN fiber populations with homogeneous myelination patterns had decreased peak amplitude and increased latency to the peak when L_u_ was longer than the putative normal length of 10 μm (Fig 5A) and L_h_ was longer than the putative normal length of 1 μm (Fig 6A). The amplitude decrease was highly significant for L_u_ > 12 μm and L_h_ > 3 μm with ~80% of a drop from normal without or with including recruitment of additional fibers with increasing SPL (Figs 5B, C and 6B, C, respectively). This was due to the fact that at those values failure of spike generation occured because of the increased lengths, L_u_ and L_h_. CAP peak latencies were significantly longer than normal for all homogeneous populations, with L_u_ > 12 μm and L_h_ > 3 μm having ~40% of an increase. The changes in CAP widths were minimal for all cases, and only significant when L_u_=13 μm or L_u_=14 μm and L_h_=4 μm or L_h_=5 μm with additional fiber recruitment (Figs 5C and 6C). For populations with heterogeneous myelination patterns, however, CAP peaks were significantly (~50%) lower, and latencies and widths were significantly higher than normal populations.

**Fig 5.**
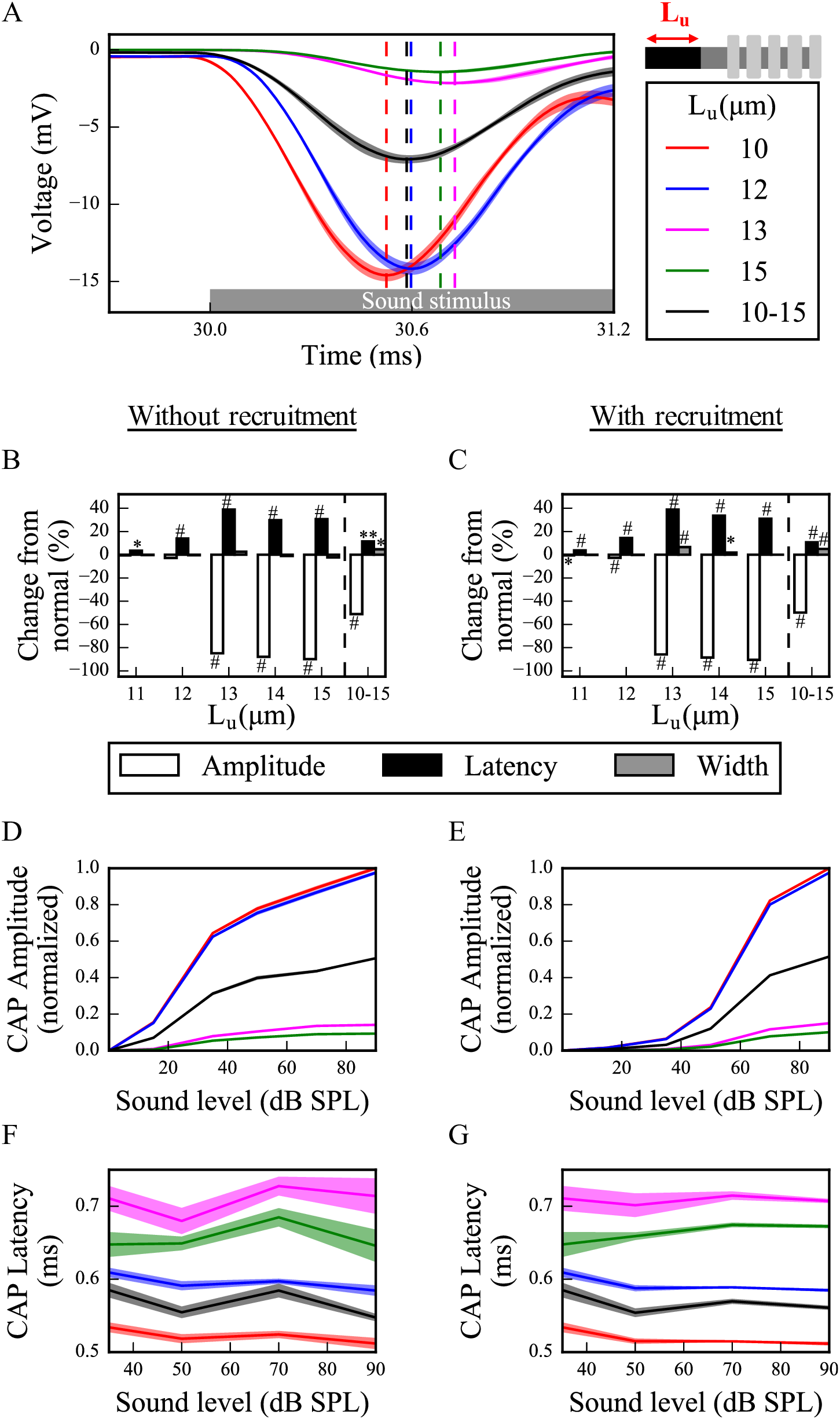
Longer L_u_ significantly decreases and delays the peak of the sound-evoked CAPs of SGN fibers. (A) Sound-evoked CAPs of SGN fiber populations with varying L_u_ at 70dB SPL without recruitment of additional fibers, averaged over 50 simulations. Shaded regions correspond to the standard error of the mean and dashed lines correspond to the peaks of each CAP, labeled with the same colors as the CAPs. The decrease and delay of peak CAPs are more obvious for populations with L_u_ > 12 μm. Comparison of CAP measures of each population relative to normal L_u_ (L_u_ = 10 μm) for cases without (B) and with (C) recruitment at 70 dB SPL. Latencies are significantly higher for all populations and peaks are significantly lower for populations with L_u_>12 μm. The increases in widths are only minimal, however significant for the heterogeneous population, where 10 μm ≤ L_u_ ≤ 15 μm (*p<0.05, **p<0.005, #p<0.0005). Normalized CAP amplitudes without (D) and with (E) recruitment for various sound levels show qualitatively similar behavior, but the case with recruitment (E) exhibits a more exponential increase. The latencies of CAP peaks increase with higher L_u_ for all sound levels with no change along the sound levels, and similar values without (F) and with (G) recruitment.

**Fig 6.**
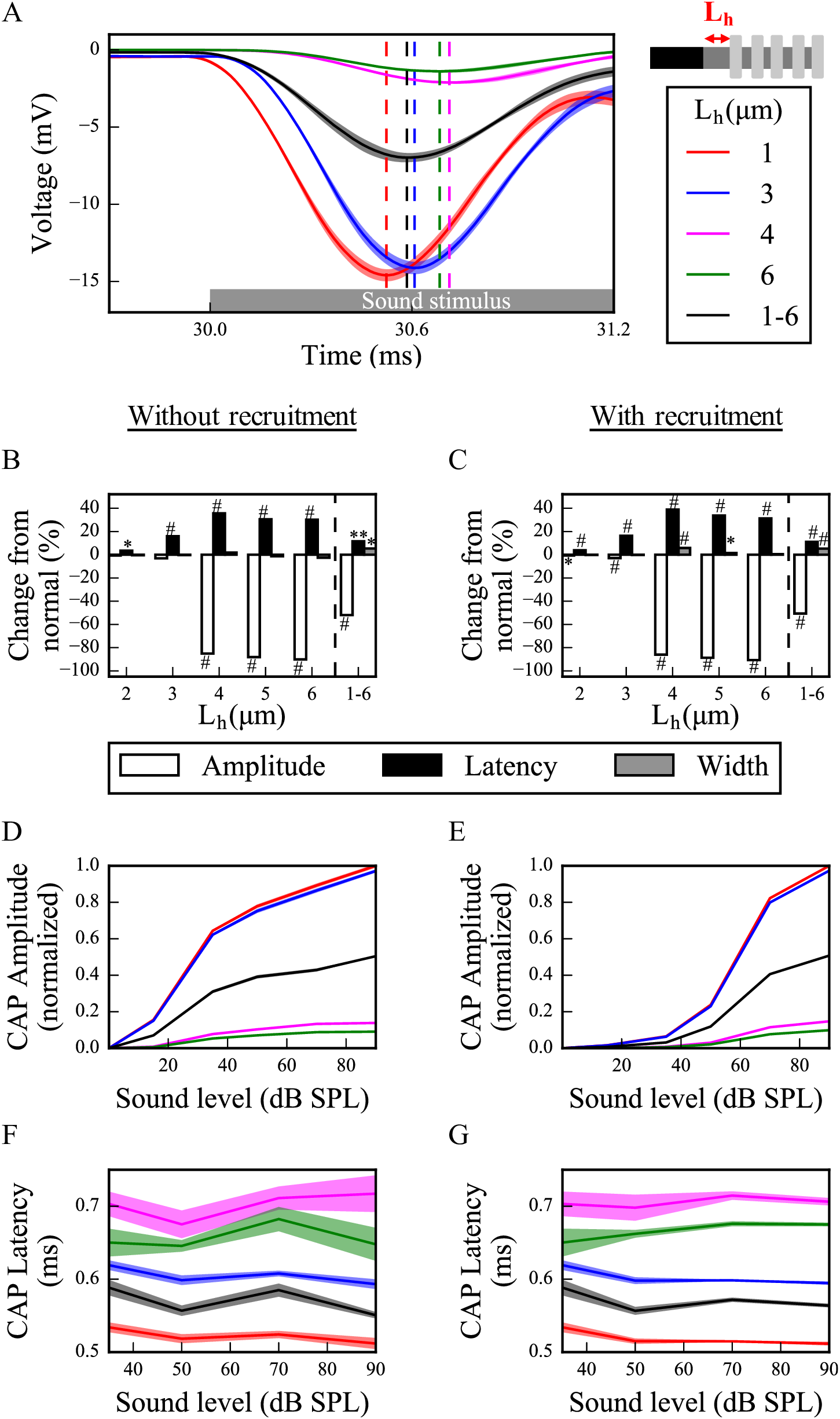
Longer L_h_ significantly decreases and delays the peak of the sound-evoked CAPs of SGN fibers. (A) Sound-evoked CAPs of SGN fiber populations of varying L_h_ at 70dB SPL without recruitment of additional fibers, averaged over 50 simulations. Shaded regions correspond to the standard error of the mean and dashed lines correspond to the peaks of each CAP, labeled with the same colors as the CAPs. The decreased peak amplitude and increased latency of CAP peak are more obvious for populations with L_h_ > 3 μm. Comparison of CAP measures of each population relative to the normal L_h_ (L_h_ = 1 μm) without (B) and with (C) recruitment at 70 dB SPL. CAP latencies are significantly higher for all populations and peak amplitudes are significantly lower for populations with L_h_>3 μm. The increases in widths are only minimal, however significant for the heterogeneous population, where 1 μm ≤ L_h_ ≤ 6 μm (*p<0.05, **p<0.005, #p<0.0005). Normalized CAP amplitudes without (D) and with (E) recruitment for various sound levels show qualitatively similar behavior, but the case with recruitment (E) exhibits a more exponential increase. The latencies of CAP peaks increase with higher L_h_ for all sound levels with no change along the sound levels, both without (F) and with (G) recruitment.

In addition, to assess the dependencies of CAP properties on sound intensities, we measured responses to simulated sound stimuli between 0-90 dB SPL with and without additional fiber recruitment. For L_u_ ≤ 12 μm and L_h_ ≤ 3 μm, CAP peak amplitudes increased with sound intensity (Figs 5D and 6D, respectively) and when fiber recruitment was included (Figs 5E and 6E, respectively) the profile of increase was more similar to experimental measurements (see Supplementary Fig 4 in [6]). However, for L_u_ > 12 μm and L_h_ > 3 μm, CAP amplitudes remained small for all sound intensities, with and without recruitment, due to reduced spike generation. For populations with heterogeneous myelination patterns, CAP amplitudes were between the L_u_=12 μm and L_u_=13 μm cases, and the L_h_=3 μm and L_h_=4 μm cases for all sound levels, reflecting reduced spike generation in some fibers of the population with higher L_u_ and L_h_ values. CAP latencies were longer for higher values of L_u_ and L_h_ with (Figs 5G and 6G) or without (Figs 5F and 6F) recruitment, but did not exhibit significant changes with varying sound level. In the heterogeneous populations, CAP latencies showed values between the L_u_=10 μm and L_u_=12 μm cases and the L_h_=1 μm and L_h_=3 μm cases.

### Effects of synaptopathy on SGN population activation patterns

There is strong evidence indicating that noise-induced synaptopathy, primarily at HT fibers, is one of the mechanisms of hidden hearing loss [8]. To simulate it, we considered responses of a population of control SGN fibers (L_u_ = 10 μm, L_h_ = 1 μm) with 50% of HT IHC-SGN synapses removed. To investigate the specific effect of loss of synapses on HT fibers, we compared responses to the case where the same number of synapses (1/6^th^ of whole population) were removed randomly from the whole population of three fiber types. The CAPs computed from populations with and without synaptopathy (Fig 7A) in response to a 70 dB SPL suggest that HT-targeted synaptopathy produces only a small effect on CAP peak amplitude while random synaptopathy has a much broader and significant effect on the amplitude (~80% vs ~10% decrease from normal at 70dB SPL) (Figs 7B and C). Moreover, there was no latency and width changes for HT synaptopathy with or without fiber recruitment. However, random synaptopathy significantly increased width and latency with recruitment at 70dB SPL, even though the increase was minimal (<1%).

**Fig 7.**
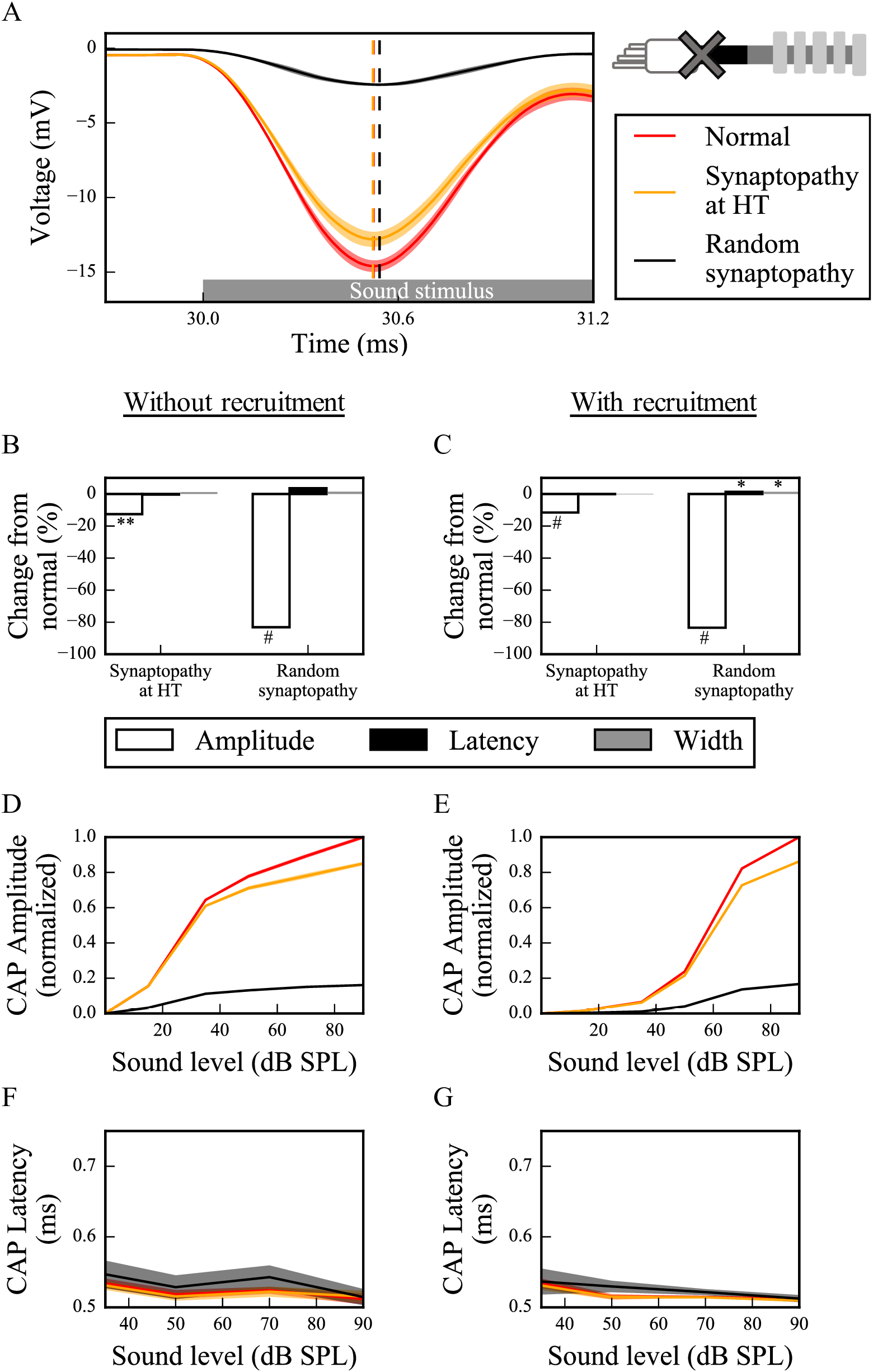
Synaptopathy at IHC-SGN synapses decreases the peak of the CAP significantly, without changes to peak latency and width. (A) Sound-evoked CAPs of SGN fiber populations with different synaptopathy scenarios at 70dB SPL without recruitment of additional fibers, averaged over 50 simulations. Shaded regions correspond to the standard error of the mean and dashed lines correspond to the peaks of each CAP, labeled with the same colors as the CAPs. Synaptopathy has smaller effects on CAP peak amplitude and latency when it affects only HT fiber synapses compared to affecting all fiber types randomly. Comparison of CAP measures of synaptopathy cases relative to normal (no synaptopathy) without (B) and with (C) recruitment at 70 dB SPL (*p<0.05, **p<0.005, #p<0.0005). Normalized CAP amplitudes show qualitatively similar behavior without (D) and with (E) recruitment for various sound levels, but the case with recruitment exhibits a more exponential increase. The latencies of the CAP peaks do not exhibit any significant difference between without (F) and with (G) recruitment cases.

We simulated sound intensities between 0-90 dB SPL with and without recruitment to assess how CAP peak amplitude and latency depend on sound intensities in the synaptopathic cochlear model. For random synaptopathy CAP peaks remained small for all sound intensities while for HT synaptopathy, a decrease of CAP peaks was observed for higher sound intensities. These results hold in cases with (Fig 7E) and without (Fig 7D) recruitment, but the profiles of CAP peak amplitudes are more consistent with experimental observations when fiber recruitment was included (see Supplementary Fig 4 in [6]). CAP latencies did not show any significant differences for any sound level in any synaptopathy case with or without recruitment (Figs 7F and G).

### Combined effects of myelinopathy and synaptopathy of hidden hearing loss

To investigate how different HHL mechanisms interact and affect cumulative SGN fiber activity, we combined them in our model (Fig 8). When HT synaptopathy (Fig 8A and B) was combined with myelinopathy affecting the length of the initial unmyelinated segment L_u_, CAP peak amplitude showed significant additive decrease but latency and width showed no change beyond that produced by the myelin defects alone (compare Case 3 with Cases 1 and 2). When both myelinopathy mechanisms were combined by varying L_u_ and L_h_ across the population, both CAP peak amplitude and latency showed significant additive changes (compare Case 4 with Case 2). CAP widths were significantly increased only by myelinopathy mechanisms, in response to simulated 70dB SPL. In response to varied sound intensities between 0-90 dB SPL, the additive effects of synaptopathy on CAP peak amplitude changes were prominent for higher SPL while latencies showed little dependence on SPL (Figs 8C and D, respectively).

**Fig 8.**
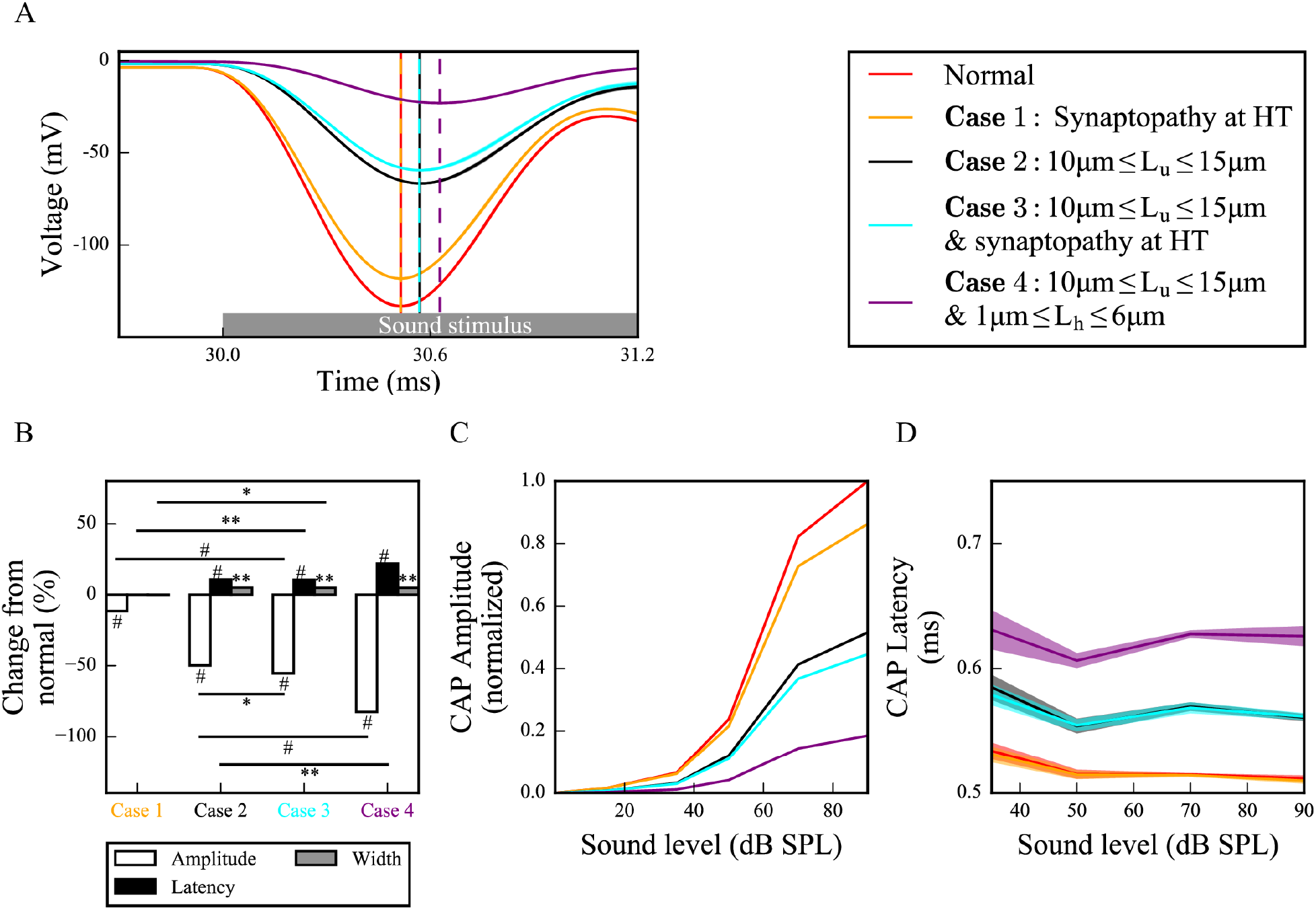
Different scenarios of hidden hearing loss have additive effects on SGN activity. (A) Sound-evoked CAPs of SGN fiber populations with different myelinopathy and synaptopathy scenarios at 70dB SPL with recruitment of additional fibers, averaged over 50 simulations (dashed lines correspond to the peaks of each CAP, labeled with the same colors as the CAPs). Combined synaptopathy and myelinopathy (Case 3) shows additive effects on the decrease in CAP peak amplitude, but not on the increase in CAP peak latency (compare to Cases 1 and 2). Combined different myelinopathies show additive effects on both CAP peak amplitude and latency (compare Cases 2 and 4). (B) Comparison of average CAP measures for different myelinopathy and synaptopathy cases relative to normal, and between cases with recruitment at 70 dB SPL (*p<0.05, **p<0.005, #p<0.0005). Normalized CAP amplitudes (C) and CAP latencies (D) for different myelinopathy and synaptopathy cases with recruitment for various sound levels, averaged over 50 simulations. Shaded areas correspond to the standard error of the mean.

In summary, model results suggest that decreases in CAP peak amplitudes show additive effects for combined synaptopathy and myelinopathy. Also, there were significant increases in CAP peak latencies and CAP widths only for myelinopathy-based mechanisms, with latencies showing additive effects in combined myelinopathies, while synaptopathies do not affect this CAP features.

## Discussion

We built a reduced biophysical model simulating sound-evoked activity of type I SGN populations to analyze two hypotheses of the cause of HHL, synaptopathy and myelinopathy. Model SGN spike times were convolved with the unitary response of the CAP, a near-field response of SGNs, to convert spike times into cumulative activity for comparison with experimental results. The model shows that synaptopathy reduces the amplitude of the cumulative CAP response without affecting its latency due to a reduction in the number of nerve fibers responding without disruption of spike timing. In contrast, myelinopathy, when modeled as disorganization of either the initial unmyelinated nerve segment length or the heminodal spacing, causes disruption of spike timing in addition to loss of firing response, affecting both the peak amplitude and latency of the cumulative CAP. Similar results are obtained when additional fibers, associated with neighboring characteristic frequencies are recruited in response to high SPL stimuli.

Previously, it has been shown that noise exposure and aging cause HHL due to synapse loss at SGN-IHC synapses, which results in a decrease of ABR P1 without increases in latency or thresholds [4]. Moreover, it has been hypothesized that synapse loss occurs preferentially at HT SGN-IHC synapses [8]. Consistent with experimental results, our simulations for both HT synaptopathy and random synaptopathy show that CAP latencies are unchanged for either scenario, but the amplitude of the CAP peak is significantly decreased. However, the decrease in CAP amplitude was much larger for random synaptopathy and no significant activity was observed for lower SPL stimuli, in contrast to HT synaptopathy case, where differences in SGN fiber activity appeared only with higher SPL stimuli. These results suggest that synaptopathy at HT synapses is a more likely scenario for HHL than random loss of synapses, since experimental results show that thresholds remain unchanged (Fig 7) [8].

As shown by Wan and Corfas (2017), myelinopathy affects the distance from the IHC-SGN synapse to the heminode and introduces heterogeneity in heminode locations across a SGN fiber population, which is likely to result in their desynchronized activity [6]. Here, we provided evidence that increasing heterogeneity of heminode locations decreases the synchronization of spike timing of SGN fiber populations. Moreover, spike rates of more heterogeneous SGN fiber populations dropped, suggesting a loss of spike generation in SGN fibers with heminodes further from IHCs (Fig 4). Our simulations of cumulative CAP signals show that myelinopathy increases the latency and the width of the peak of CAP, similar to experimental observations of ABR P1, providing support for the disruption of spike timing in SGN activity (Figs 5 and 6). In addition, the amplitude of the simulated CAP decreased with myelinopathy, reflecting the reduction of SGN spike activity.

Combining synaptopathy and myelinopathy HHL mechanisms led to additive effects in our model. Decreases in CAP peak amplitude were additive for combined synaptopathy and myelinopathy, but synaptopathy did not contribute to changes in CAP latency even in the combined scenario. Combining myelinopathy mechanisms led to additive increases in both peak CAP amplitude and latency (Fig 8). These results match with the experimental results qualitatively, further supporting the accuracy of our model.

In the myelinopathy simulations, we varied the length of the initial unmyelinated segment L_u_ keeping a constant channel density (Fig 5) and varied the length of the heminode L_h_ keeping constant channel numbers (Fig 6). Results show similar effects on SGN fiber activity, i.e. the populations with the same combined lengths L_u_+L_h_ exhibit the same behavior. As evidence on how channels might be affected by the disruption of myelination patterns is lacking, we also simulated cases where L_u_ increases with constant channel number (S1 Fig) and L_h_ increases with constant channel density (S2 Fig). Results show that spreading the same number of channels over an increased L_u_ (S1 Fig), rather than increasing the number by keeping the channel density constant (Fig 5), decreases the L_u_ value at which the abrupt decrease in CAP peak occurs due to loss of spike generation. With constant channel number, CAP peaks for homogeneous populations with L_u_>11 μm decreased ~90% from normal (L_u_=10 μm) (S1B Fig). However, the same drop occurred when L_u_>12 μm for the constant channel density case. In contrast, varying L_h_ while keeping the heminode channel density constant, i.e., increasing the number of channels for larger L_h_, increased the L_h_ value associated with the loss of spike generation up to 6 μm, compared to 3 μm when channel number was kept constant (Fig 6). To conclude, any of these scenarios results in qualitatively similar SGN fiber activity patterns, only affecting the L_u_ and L_h_ lengths at which loss of spike generation leads to an abrupt drop in the CAP peak.

To better understand the effects of myelinopathy on SGN spike generation, we additionally analyzed the population outcome of vesicle release events to the SGN fibers. As described in the Methods section, SGN response to vesicle release was simulated by applying a brief external current pulse to the peripheral end of the SGN fibers. We thus calculated the probability that release events result in corresponding spikes for various amplitudes I_app_ of the external current pulse for increasing values of L_u_ (S3A Fig). For simulated 70dB SPL stimuli, higher I_app_ amplitudes increased spike probability for larger L_u_ values, leading to increases in the L_u_ values at which spike generation was affected. If L_u_ exceeded a critical value, the probability of spike generation decreased significantly. These results show that this L_u_ critical value required for spike generation depends on IHC-SGN synaptic efficacy.

To analyze the effect of sound level on SGN fiber spike probability, we ran simulations for all sound levels keeping I_app_ fixed at the default value (I_app_ = 0.024nA, solid black rectangle in S3A Fig). As described in the Methods section, increasing sound level was simulated by increasing the probability of a vesicle release event, thus leading to higher rate of release from IHCs, i.e. higher frequencies of external current pulse applications to SGN fibers. For this I_app_ value, spike generation was affected for L_u_>12 μm as evident in the results shown in Fig 5. For SGN fibers with L_u_ ≤ 12.3 μm, spike probabilities were higher than 70% for all sound levels (S3B Fig). However, spike probabilities decreased gradually with higher sound levels due to the inability of the fibers to respond to high frequency stimulation. This means, despite more frequent release events from IHC-SGN synapses with higher sound levels, SGN fibers cannot fire with a higher frequency due to the saturation of their spike rate, resulting in decreased spike probabilities. For SGN fibers with L_u_>12.3 μm, spike probability was very low reflecting loss of spike generation but it increased slightly with increasing sound level, as high frequency stimulation facilitated spike generation due to temporal summation. Results for heterogeneous L_u_ values between 10 and 15 μm showed intermediate spike probabilities (~40%) as compared to homogeneous L_u_ values of 10 μm, for all sound levels.

Lastly, to analyze effects of myelinopathy on SGN spike latency, we averaged the time differences between each spike and the preceding release event causing the spike for populations of SGN fibers with varied homogeneous L_u_ values and varied sound levels (S3C Fig). The populations with L_u_>12 μm were not included since spikes were not reliably generated and for the heterogeneous population, the fibers with L_u_>12 μm were ignored. The homogeneous populations showed increased latencies with increasing L_u_ and the heterogeneous population’s latencies were between those for L_u_ = 11 μm and L_u_ = 12 μm. Latencies showed little dependence on sound levels. However, standard deviations of spike latencies increased with sound level, presumably reflecting higher variability in spike response to higher frequency stimulation (S3D Fig). Additionally, the population with heterogeneous L_u_ values showed higher standard deviations for all sound levels than the homogeneous populations with L_u_ ≤ 12 μm. This increase in spike timing variability is responsible for increases in the width of the cumulative CAP for the heterogeneous population shown in Fig 5.

In conclusion, our model results show that HHL deficits due to myelinopathy could be caused by not only loss of SGN spike activity, as in synaptopathy, but also disruption of spike timing and synchronization across a population of SGN fibers. Illumination of the underlying differences in these mechanisms for HHL based on the model may be useful for the development and testing of treatments for HHL. Moreover, the model framework may be extended to investigate mechanisms behind other peripheral auditory system disorders.

## Notes

### Competing Interest Statement

The authors have declared no competing interest.

## References

1. Kohrman DC, Wan G, Cassinotti L, Corfas G. Hidden Hearing Loss: A Disorder with Multiple Etiologies and Mechanisms. Cold Spring Harb Perspect Med. 2019. Epub 2019/01/07. doi: 10.1101/cshperspect.a035493. PubMed PMID: 30617057.

2. Spankovich C, Gonzalez VB, Su D, Bishop CE. Self reported hearing difficulty, tinnitus, and normal audiometric thresholds, the National Health and Nutrition Examination Survey 1999-2002. Hear Res. 2017. Epub 2017/12/07. doi: 10.1016/j.heares.2017.12.001. PubMed PMID: 29254853.

3. Tremblay KL, Pinto A, Fischer ME, Klein BE, Klein R, Levy S, et al. Self-Reported Hearing Difficulties Among Adults With Normal Audiograms: The Beaver Dam Offspring Study. Ear Hear. 2015;36(6):e290–9. doi: 10.1097/AUD.0000000000000195. PubMed PMID: 26164105; PubMed Central PMCID: PMCPMC4824300.

4. Kujawa SG, Liberman MC. Adding insult to injury: cochlear nerve degeneration after “temporary” noise-induced hearing loss. J Neurosci. 2009;29(45):14077–85. doi: 10.1523/JNEUROSCI.2845-09.2009. PubMed PMID: 19906956; PubMed Central PMCID: PMCPMC2812055.

5. Sergeyenko Y, Lall K, Liberman MC, Kujawa SG. Age-related cochlear synaptopathy: an early-onset contributor to auditory functional decline. J Neurosci. 2013;33(34):13686–94. doi: 10.1523/JNEUROSCI.1783-13.2013. PubMed PMID: 23966690; PubMed Central PMCID: PMCPMC3755715.

6. Wan G, Corfas G. Transient auditory nerve demyelination as a new mechanism for hidden hearing loss. Nat Commun. 2017;8:14487. Epub 2017/02/17. doi: 10.1038/ncomms14487. PubMed PMID: 28211470; PubMed Central PMCID: PMCPMC5321746.

7. Rajan R, Cainer KE. Ageing without hearing loss or cognitive impairment causes a decrease in speech intelligibility only in informational maskers. Neuroscience. 2008;154(2):784–95. Epub 2008/04/04. doi: 10.1016/j.neuroscience.2008.03.067. PubMed PMID: 18485606.

8. Furman AC, Kujawa SG, Liberman MC. Noise-induced cochlear neuropathy is selective for fibers with low spontaneous rates. J Neurophysiol. 2013;110(3):577–86. Epub 2013/04/17. doi: 10.1152/jn.00164.2013. PubMed PMID: 23596328; PubMed Central PMCID: PMCPMC3742994.

9. Long P, Wan G, Roberts MT, Corfas G. Myelin development, plasticity, and pathology in the auditory system. Dev Neurobiol. 2018;78(2):80–92. Epub 2017/09/26. doi: 10.1002/dneu.22538. PubMed PMID: 28925106; PubMed Central PMCID: PMCPMC5773349.

10. Takazawa T, Ikeda K, Murata K, Kawase Y, Hirayama T, Ohtsu M, et al. Sudden deafness and facial diplegia in Guillain-Barré Syndrome: radiological depiction of facial and acoustic nerve lesions. Intern Med. 2012;51(17):2433–7. Epub 2012/09/01. PubMed PMID: 22975563.

11. Choi JE, Seok JM, Ahn J, Ji YS, Lee KM, Hong SH, et al. Hidden hearing loss in patients with Charcot-Marie-Tooth disease type 1A. Sci Rep. 2018;8(1):10335. Epub 2018/07/09. doi: 10.1038/s41598-018-28501-y. PubMed PMID: 29985472; PubMed Central PMCID: PMCPMC6037750.

12. Kiang NY, Rho JM, Northrop CC, Liberman MC, Ryugo DK. Hair-cell innervation by spiral ganglion cells in adult cats. Science. 1982;217(4555):175–7. PubMed PMID: 7089553.

13. Hines ML, Carnevale NT. NEURON: a tool for neuroscientists. Neuroscientist. 2001;7(2):123–35. doi: 10.1177/107385840100700207. PubMed PMID: 11496923.

14. Mino H, Rubinstein JT, Miller CA, Abbas PJ. Effects of electrode-to-fiber distance on temporal neural response with electrical stimulation. IEEE Trans Biomed Eng. 2004;51(1):13–20. doi: 10.1109/TBME.2003.820383. PubMed PMID: 14723489.

15. Møller AR, Colletti V, Fiorino FG. Neural conduction velocity of the human auditory nerve: bipolar recordings from the exposed intracranial portion of the eighth nerve during vestibular nerve section. Electroencephalogr Clin Neurophysiol. 1994;92(4):316–20. doi: 10.1016/0168-5597(94)90099-x. PubMed PMID: 7517853.

16. Woo J, Miller CA, Abbas PJ. The dependence of auditory nerve rate adaptation on electric stimulus parameters, electrode position, and fiber diameter: a computer model study. J Assoc Res Otolaryngol. 2010;11(2):283–96. Epub 2009/12/22. doi: 10.1007/s10162-009-0199-2. PubMed PMID: 20033248; PubMed Central PMCID: PMCPMC2862915.

17. Woo J, Miller CA, Abbas PJ. Biophysical model of an auditory nerve fiber with a novel adaptation component. IEEE Trans Biomed Eng. 2009;56(9):2177–80. Epub 2009/06/02. doi: 10.1109/TBME.2009.2023978. PubMed PMID: 19497810.

18. Glowatzki E, Fuchs PA. Transmitter release at the hair cell ribbon synapse. Nat Neurosci. 2002;5(2):147–54. doi: 10.1038/nn796. PubMed PMID: 11802170.

19. Winter IM, Robertson D, Yates GK. Diversity of characteristic frequency rate-intensity functions in guinea pig auditory nerve fibres. Hear Res. 1990;45(3):191–202. PubMed PMID: 2358413.

20. Smith RL, Zwislocki JJ. Short-term adaptation and incremental responses of single auditory-nerve fibers. Biol Cybern. 1975;17(3):169–82. PubMed PMID: 1125344.

21. Keithley EM, Schreiber RC. Frequency map of the spiral ganglion in the cat. J Acoust Soc Am. 1987;81(4):1036–42. PubMed PMID: 3571719.

22. Liberman MC. Auditory-nerve response from cats raised in a low-noise chamber. J Acoust Soc Am. 1978;63(2):442–55. PubMed PMID: 670542.

23. Louage DH, van der Heijden M, Joris PX. Temporal properties of responses to broadband noise in the auditory nerve. J Neurophysiol. 2004;91(5):2051–65. doi: 10.1152/jn.00816.2003. PubMed PMID: 15069097.

24. Bourien J, Tang Y, Batrel C, Huet A, Lenoir M, Ladrech S, et al. Contribution of auditory nerve fibers to compound action potential of the auditory nerve. J Neurophysiol. 2014;112(5):1025–39. Epub 2014/05/21. doi: 10.1152/jn.00738.2013. PubMed PMID: 24848461.

