## Supplementary material for "Contrasting mechanisms for hidden hearing loss: synaptopathy vs myelin defects": S1 Fig

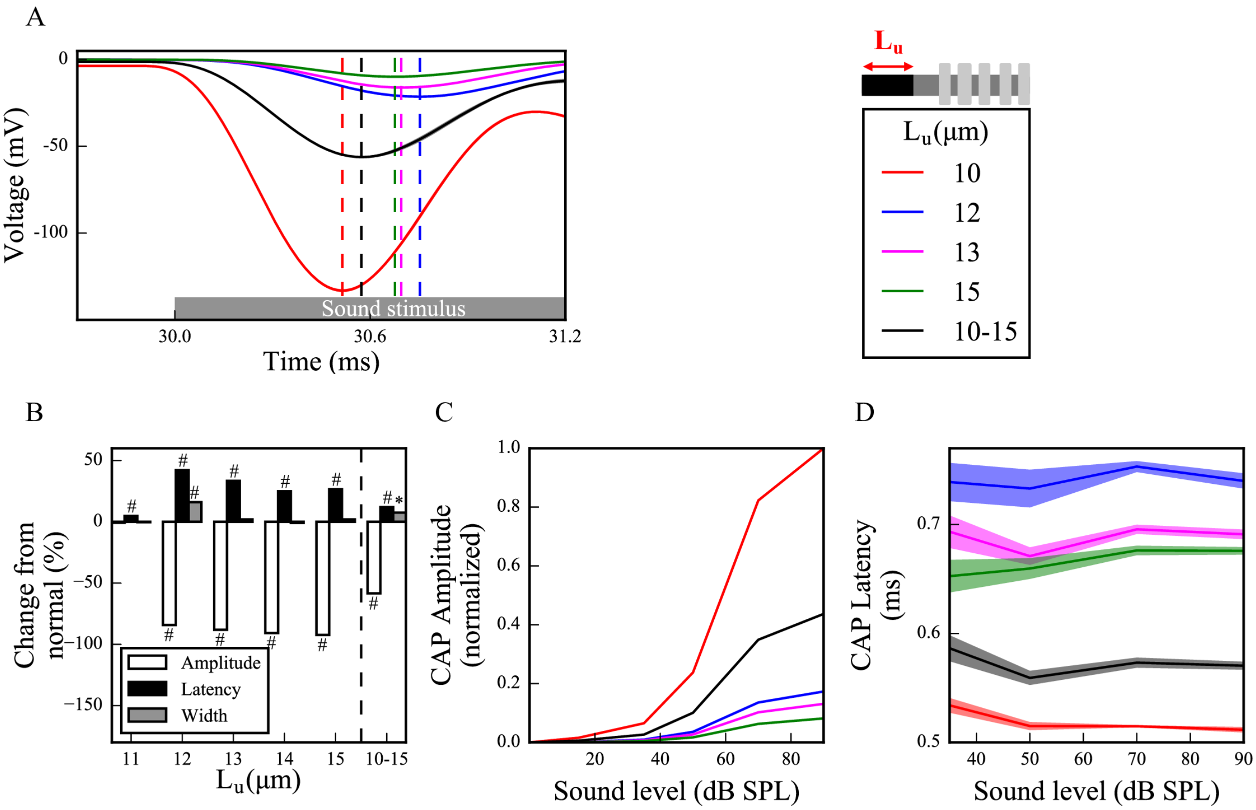


**S1 Fig. Keeping constant channel number as length of unmyelinated segment, $L_u$, is increased leads to larger effects on cumulative CAP of increased $L_u$.**

(A) Sound-evoked CAPs of SGN fiber populations with varied $L_u$ at 70dB SPL including recruitment of additional fibers, averaged over 50 simulations (dashed lines correspond to the peaks of each CAP, labeled with the same colors as the CAPs). The number of membrane ionic channels was kept fixed at the values for normal $L_u$ ($L_u$ =10 μm). Decreases in peak amplitude and increases in peak latency are larger for populations with $L_u > 11$ μm (compare to Fig 5). (B) Comparison of CAP measures relative to normal $L_u$ ($L_u$ =10 μm) of each population with recruitment at 70 dB SPL (*$p<0.05$, **$p<0.005$, #$p<0.0005$). Normalized CAP amplitudes (C) and CAP latencies (D) with recruitment for various sound levels, averaged over 50 simulations. Shaded areas correspond to the standard error of the mean.
