## Supplementary material for "Contrasting mechanisms for hidden hearing loss: synaptopathy vs myelin defects": S1 File

**S1 File. Model equations.**

For each SGN fiber, the transmembrane potential $V_m$ is a function of space x and time t and is expressed as:

$$-\frac{1}{R_a}\frac{\partial^2 V_m(x,t)}{\partial x^2} + C_m\frac{\partial V_m(x,t)}{\partial t} + \frac{V_m(x,t)-E_{rest}}{R_m} + I_{ion}(x,t) = I_{app}(x,t) \quad (1)$$

where $R_a$ is the specific cytoplasmic resistance, $C_m$ is the specific capacitance, $R_m$ is the specific membrane resistance, $E_{rest}$ is the resting potential, $I_{ion}(x,t)$ and $I_{app}(x,t)$ are ionic and applied currents, respectively.

Ionic current ($I_{ion}(x,t)$) consists of sodium ($I_{Na}(x,t)$) and potassium ($I_K(x,t)$) currents:

$$I_{ion}(x,t) = I_{Na}(x,t) + I_K(x,t) \quad (2)$$

where,

$$I_{Na}(x,t) = g_{Na}\big(m(t)\big)^3 h(t)(V_m(x,t) - E_{Na}) \quad (3)$$

and

$$I_K(x,t) = g_K\big(n(t)\big)^4 (V_m(x,t) - E_K) \quad (4)$$

Here, m(t), h(t) and n(t) are gating variables, $g_{Na}$ and $g_K$ are maximal sodium and potassium conductances, respectively, and $E_{Na}$ and $E_K$ are the Nernst potentials for sodium and potassium ions, respectively. The gating variables i (for i=m, n and h) are expressed in terms of rate functions $\alpha_i(V_m)$ and $\beta_i(V_m)$, such that [14]:

$$\frac{di}{dt} = \alpha_i(V_m)(1-i) - \beta_i(V_m)i \;\; for\; i = m,n,h \quad (5)$$

where,

$$\alpha_m(V_m) = \frac{1.872(V_m+52.59)}{1-e^{\frac{-(V_m+52.59)}{6.06}}} \quad (6)$$

$$\beta_m(V_m) = \frac{-3.973(V_m+57)}{1-e^{\frac{V_m+57}{9.41}}} \quad (7)$$

$$\alpha_h(V_m) = \frac{-0.549(V_m+105.74)}{1-e^{\frac{V_m+105.74}{9.06}}} \quad (8)$$

$$\beta_h(V_m) = \frac{22.57}{1+e^{\frac{-(V_m+22)}{12.5}}} \quad (9)$$

$$\alpha_n(V_m) = \frac{0.129(V_m+43)}{1-e^{\frac{-(V_m+43)}{10}}} \quad (10)$$

$$\beta_n(V_m) = \frac{-0.324(V_m+68)}{1-e^{\frac{V_m+68}{10}}} \quad (11)$$
