## Supplementary material for "Contrasting mechanisms for hidden hearing loss: synaptopathy vs myelin defects": S2 Fig

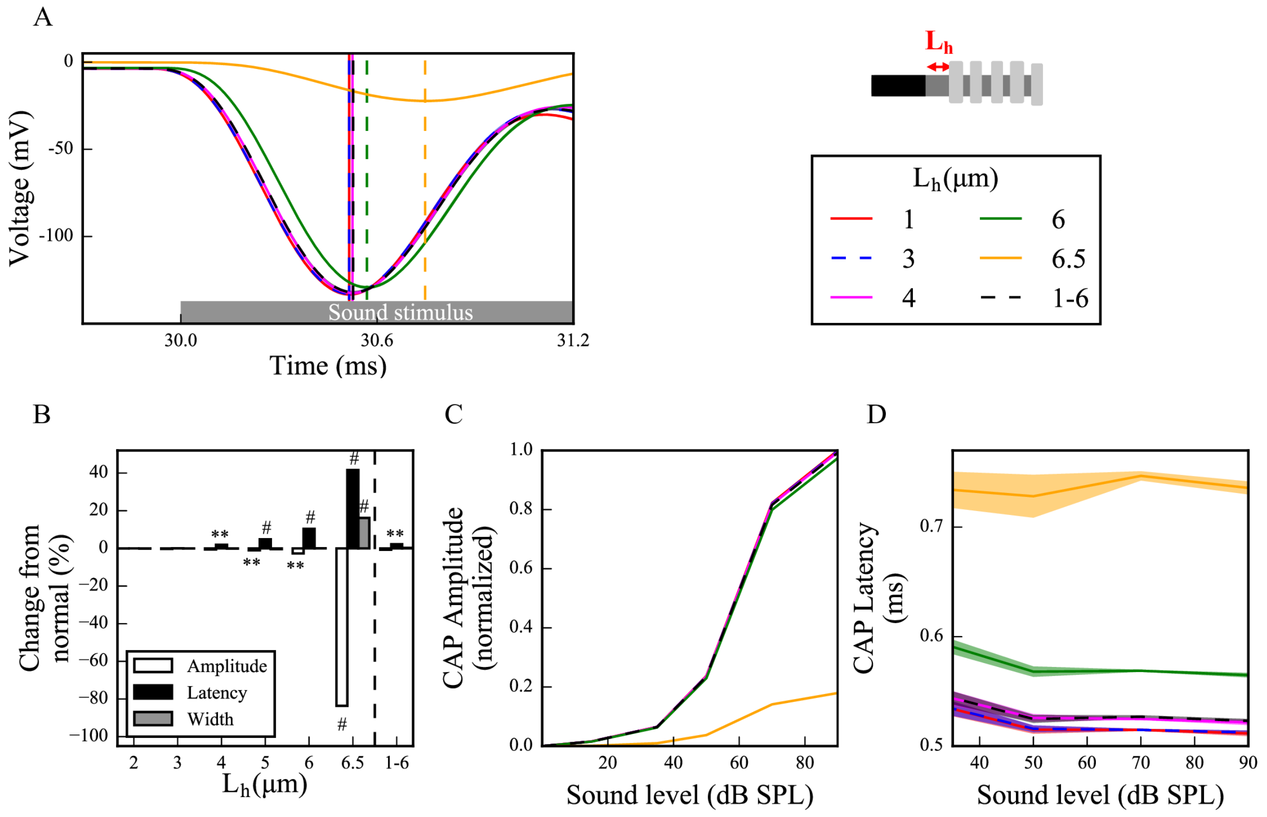


**S2 Fig. Maintaining channel density at the heminode as its length, $L_h$, is varied reduces effects on cumulative CAP of increased $L_h$.**

(A) Sound-evoked CAPs of SGN fiber populations with varied $L_h$ at 70dB SPL including recruitment of additional fibers, averaged over 50 simulations (dashed lines correspond to the peaks of each CAP, labeled with the same colors as the CAPs). Densities of membrane ionic channels were kept constant at the values for normal $L_h$ ($L_h$ =1 μm). Decreases in amplitude and increases in latency of CAP peaks are more obvious for populations with $L_h > 6$ μm (compare to Fig 6). (B) Comparison of CAP measures relative to normal $L_h$ ($L_h$ =1 μm) for each population with recruitment included at 70 dB SPL (*$p<0.05$, **$p<0.005$, #$p<0.0005$). Normalized CAP amplitudes (C) and CAP latencies (D) with recruitment for various sound levels, averaged over 50 simulations. Shaded areas correspond to the standard error of the mean.
