## Supplementary material for "Contrasting mechanisms for hidden hearing loss: synaptopathy vs myelin defects": S3 Fig

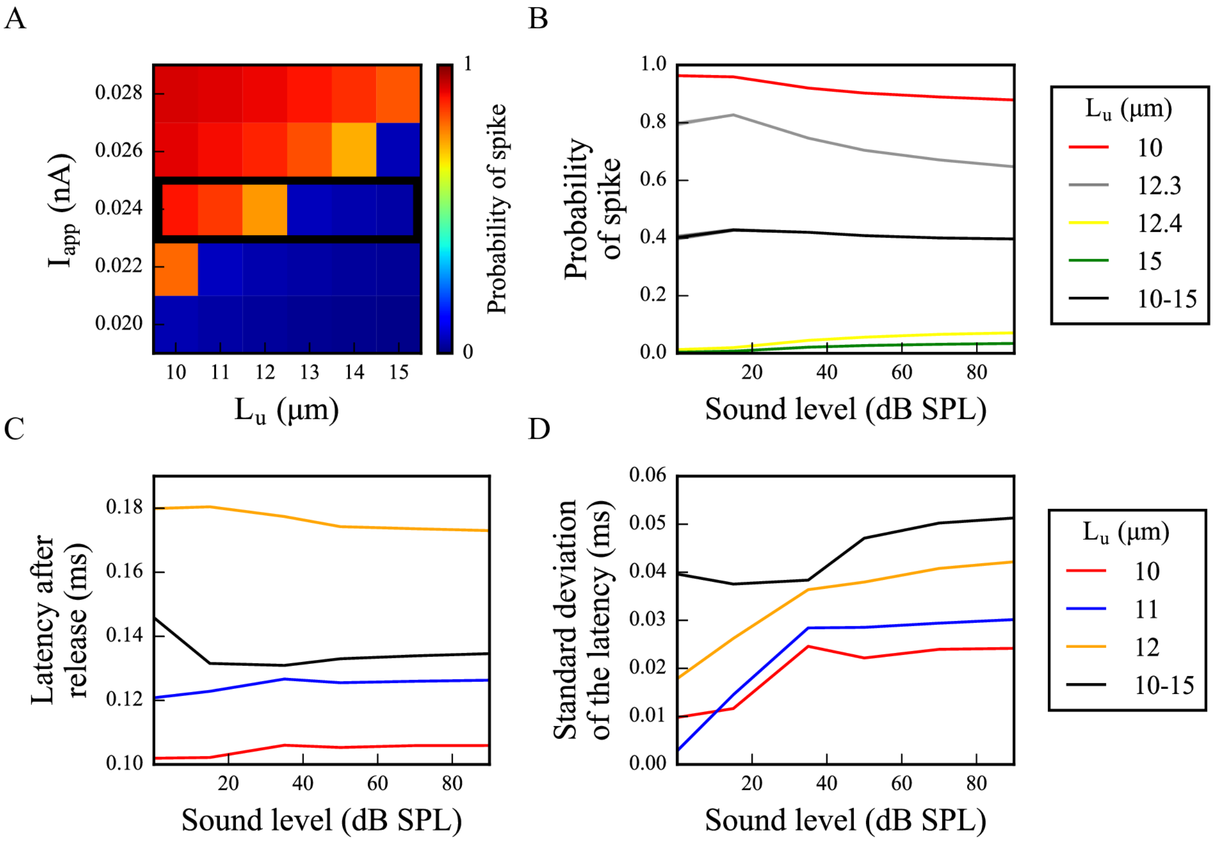


**S3 Fig. Myelinopathy results in a significantly reduced spike probability and increased latency after a release event.**

(A) The probability that simulated IHC-SGN synaptic vesicle release events result in spike generation at the heminodes of postsynaptic SGN fibers was calculated for various SGN fiber populations at 70dB SPL, averaged over 50 simulations. The amplitude of external current pulses ($I_{app}$) applied at the beginning of $L_u$, representing IHC-SGN vesicle release, was varied between 0.020nA and 0.028nA. The threshold $L_u$, where abrupt drop of spike probability occurs, increases with increasing $I_{app}$. Panels (B)-(D) and all other results in the paper were obtained with $I_{app}$=0.024nA. (B) Spike probabilities for SGN fiber populations with different homogeneous $L_u$ values in response to different sound levels exhibit an abrupt drop when $L_u \geq 12.3$ μm for all sound levels. (C) The average latency after each release event of spikes across SGN fiber populations, averaged over 50 simulations, increases for longer $L_u$. (D) Standard deviations of spike latencies of SGN fiber populations, averaged over 50 simulations, increase with sound level. The heterogeneous population (10 μm $\leq L_u \leq$ 15 μm) has higher standard deviation than homogeneous populations for every sound level. Since fibers with $L_u$ >12 μm do not fire in response to single release events, they are not shown in panels (C) and (D).
